# The longer the better: longer acrosomes contain more proteins involved in sperm-egg interactions

**DOI:** 10.1101/2023.07.04.547644

**Authors:** Tereza Otčenášková, Romana Stopková, Pavel Stopka

**Author notes:** These authors jointly supervised this work: Romana Stopková and Pavel Stopka.

## Abstract

Speciation and sperm competition have been shown to be the major driving forces for sperm morphology variation, swimming velocity and metabolism. We used light microscopy to measure sperm traits and nLC-MS/MS to detect proteomic variation in three species of rodents – promiscuous *Apodemus flavicollis,* less promiscuous *Microtus arvalis,* and the least promiscuous *Mus musculus musculus*. We show that the length of sperm apical hook containing the acrosome is the most variable trait and that this variation is reflected by proteomes on interspecific and intraspecific levels. Thus, we provide potential markers of selection such as Ldhc (in Mus and Apodemus) for long acrosomes which is a gene coding L-lactate dehydrogenase that is involved in sperm motility and Spaca1, which is important in sperm-oocyte fusion), and e.g. Mup17 for short acrosomes – a gene coding a Major urinary protein that likely chelates lipophilic compounds after spermiogenesis. In short, longer acrosomes are characteristic of proteins involved in fertilisation and gluconeogenesis, while shorter acrosomes contain more cytoskeletal proteins important for spermiogenesis. For the first time, we demonstrate that there is an innate and evolvable variability in sperm morphology and corresponding proteomes within species that can be driven by sperm competition to species-specific reproductive optima.

**Summary statement:** We show that interspecific and intraspecific variation in sperm morphology traits are detectable on proteomic level thus providing markers of selection due to sperm competition.

## 1. Introduction

Sperm competition, one of the mechanisms of postcopulatory sexual selection, has been demonstrated as a selective force that drives the evolution of reproductive traits such as reproductive behaviour, reproductive tract morphology and sperm design across many taxa including mammals (Briskie and Montgomerie, 1992, Parker, 1970, Roldan et al., 1992, Stockley et al., 1997). Diverse degrees of female promiscuity are in males reflected in interspecific variation in the ratio of testis mass corrected for body mass and thus in the amount of sperm produced (Firman and Simmons, 2008, Tourmente et al., 2011). In mammals, levels of sperm competition are positively correlated with sperm size and shape. An extensive study conducted on 226 species of Eutherian mammals revealed a positive relationship between the lengths of six sperm parameters and relative testes mass (midpiece length correlated non-significantly)(Tourmente et al., 2011). However, Gomez Montoto et al. 2011 (Gómez Montoto et al., 2011) showed that only the dimensions of the sperm head (i.e., sperm head length and area) are significantly related to the risk of sperm competition in muroid species. These findings are in contrast to passerine birds, where sperm head length was found to be a sperm trait not associated with sperm competition at an interspecific level (Støstad et al., 2018).

Furthermore, spermatozoa of muroid species with promiscuous mating strategy have longer and more reflected apical hooks with a decreasing between-male variance in the hook length (Immler et al., 2007, Šandera et al., 2013). The apical hook is a characteristic feature in murids determining the falciform shape of the sperm head but see (Breed, 2004, Gómez Montoto et al., 2011); that is unique among Eutherian mammals. In the highly promiscuous European wood mouse (*Apodemus sylvaticus*) and deer mouse (*Peromyscus maniculatas*), apical hooks were reported to mediate collective sperm behaviour via aggregation of numerous sperm into ‘sperm trains’ of higher swimming speed than individual spermatozoa (Fisher and Hoekstra, 2010, Moore et al., 2002). Nevertheless, as supported by recent cross-species studies, the main significance of apical hooks is rather in sperm interactions with the female genital tract and/or the oocyte with train formation being a trait specific to only a few species (Hook et al., 2021, Tourmente et al., 2016). Spermatozoa of muroid rodents undergo premature (also termed spontaneous) acrosome reaction that was identified as a physiological reproductive strategy (Clift et al., 2009, Johnson et al., 2007). Under competitive conditions, this ability is associated with sperm aggregates dissociation and thus faster fertilisation (Moore et al., 2002).

The risk of sperm competition is also associated with sperm performance in rodents. Unlike in songbirds (Rowe et al., 2013), spermatozoa from muroid species experiencing enhanced levels of sperm competition are of higher ATP content that results in higher swimming velocities (Tourmente et al., 2015b). Regarding the metabolic pathway for sperm energy production, mitochondria-related oxidative phosphorylation is favoured over glycolysis in more promiscuous species as observed between closely related species from the genus *Mus* (Tourmente et al., 2015a). Enhanced energy metabolism is, however, linked with a risk of higher production of reactive oxygen species. To prevent spermatozoa from potential oxidative stress damage, the fatty-acid composition of sperm membranes is altered by increasing the percentage of peroxidation-resistant fatty acids under competitive conditions (delBarco-Trillo et al., 2015). Besides sperm motility and energetics, acrosomal proteins involved in sperm-oocyte binding and fusion are under positive selection in species with high sperm competition levels (Vicens et al., 2017).

As indicated above, the majority of studies conducted on rodents describe the effects of sperm competition on sperm design, function and composition at the species level. Our study, therefore, examines whether the interspecific variation of sperm traits caused by promiscuous mating can be observed also between individuals from one species. For this purpose, three species from the muroid rodent families *Muridae* and *Cricetidae* were chosen, i.e., the house mouse (*Mus musculus musculus* - MM), the yellow-necked mouse (*Apodemus flavicollis* - AF) and the common vole (*Microtus arvalis* - MA). These species experience low, high and low-moderate levels of sperm competition, respectively, as expressed by the relative testes mass in (Tourmente et al., 2011). Based on measurements of sperm dimensions and to address proteins involved in fertilisation, we performed proteomic analysis using nano-scale liquid chromatography tandem mass spectrometry (nLC-MS/MS) of acrosomal proteins isolated from spermatozoa and we analysed two subsets of individuals with short and long apical hooks and thus acrosomes per species. Our working hypothesis assumes that spermatozoa with longer apical hooks have longer acrosomes that are similarly located in the apical region of the sperm head. The workflow is summarized in Fig. 1.

**Fig. 1.**
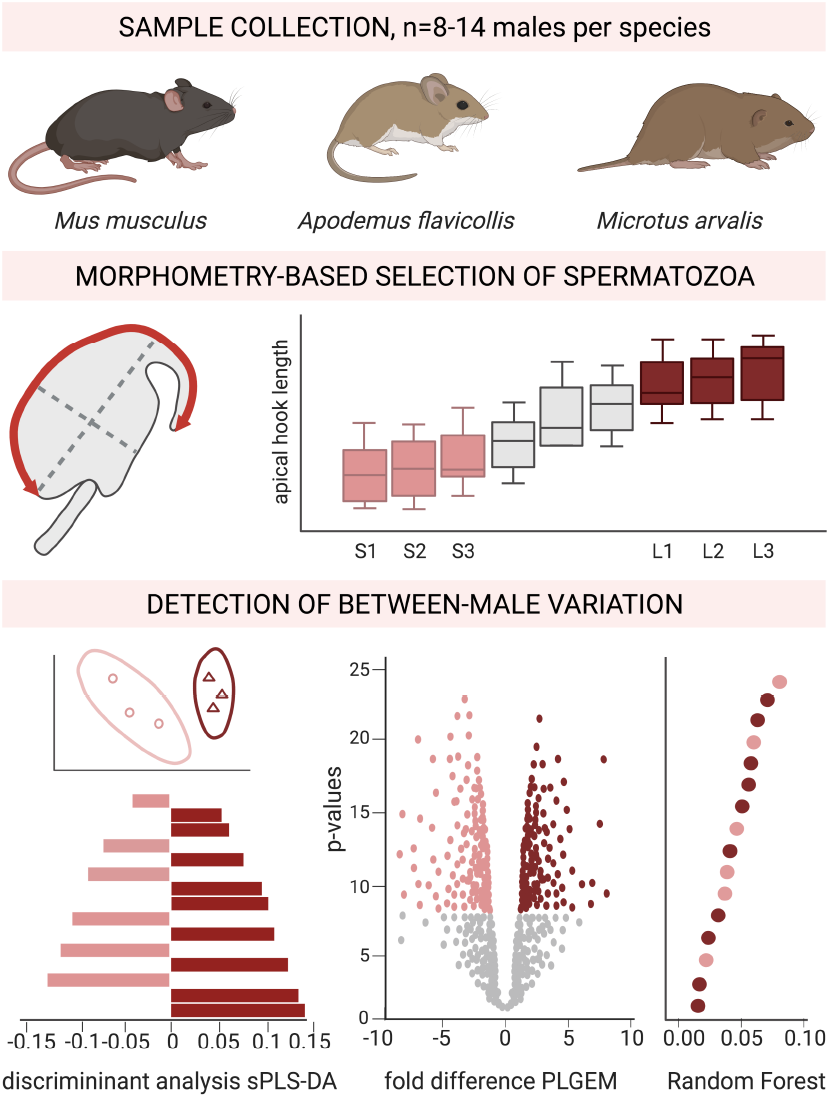
Graphical abstract. Workflow summary was created with BioRender.com.

## 2. Results

### 2.1. Proteomes of the acrosomal contents

As shown by immunofluorescent staining, spermatozoa of all three rodent species were characterized by an apical hook of species-specific shape and size, Fig. 2A. Furthermore, the length of the apical hook was the most variable sperm trait in our morphometry analysis with the following range of mean values: 12.899 μm (s.d. 0.721) - 14.113 μm (s.d. 1.146) in MM, 17.562 μm (s.d. 1.351) – 19.784 μm (s.d. 0.812) in AF and 10.541 μm (s.d. 0.934) – 12.897 μm (s.d. 1.352) in MA, Fig. 2B. Thus, we chose apical hook length as a parameter for sample selection and performed nLC-MS/MS proteomic analysis of acrosomal content isolated from three individuals with the shortest and three longest apical hooks per species. With this approach, we omitted low variation to better follow the individual differences within species. The nLC-MS/MS analysis of 18 samples (six samples per species) yielded a total of 648 protein identifications (IDs). The proteomic dataset was LFQ normalised to reduce variation of nonbiological origin (i.e., introduced during MS runs) followed by removal of incomplete rows (those with less than two IDs across all samples), identified but not quantified proteins and potential contaminants. Before further analyses, the resulting dataset containing a total of 277 IDs was Quantile-normalised to remove variation between samples. The most protein IDs (N = 239) were detected in the acrosomal proteome of AF followed by MA and MM with 194 and 179 IDs, respectively, with nearly half of all identified proteins (N = 122, 44 %) being common to all species. To explore the proteomic basis of morphological sperm variation, we provided a factor to the dataset dividing proteins detected in sperm with short apical hooks and those in sperm with long apical hooks. Inspection of data distribution revealed a similar structure of negative-binomial distribution of acrosomal proteomes across species, Fig. 2C.

**Fig. 2.**
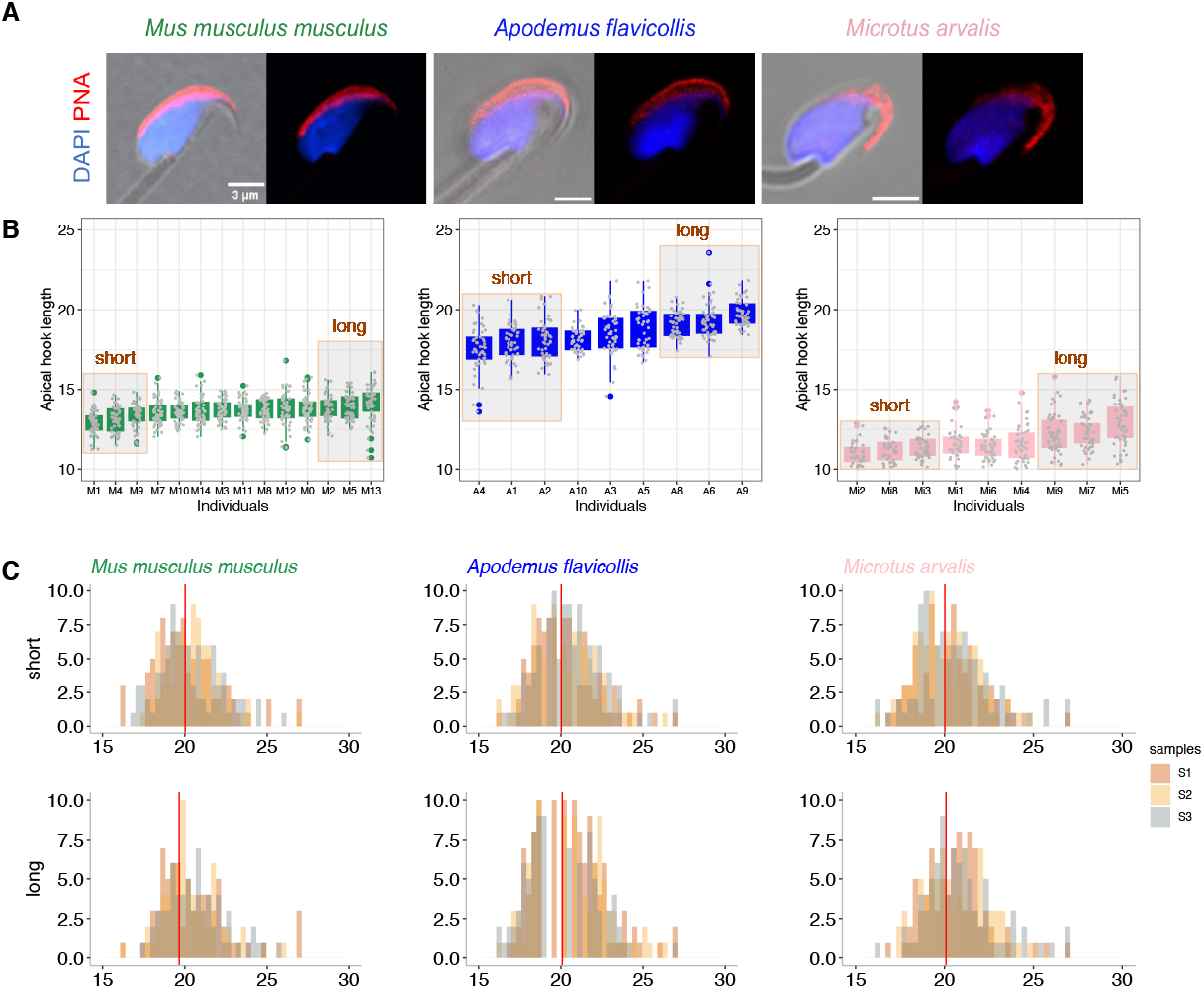
Morphometric variation and proteomes inspection. (A) Sperm head morphology in selected rodent species. Note the interspecific variation in the shape of apical hooks. Acrosomes were highlighted with PNA (red), nuclei counterstained with DAPI (blue). Left panels are merged with DIC. Scale bar, 3 μm. (B) Individual variation in the apical hook length. Only individuals with shortest and longest apical hooks were selected for further analyses, see boxed areas. A total of 50 sperm per species were measured. Measures (y-axis) are in μm. (C) Visualization of protein signal distributions revealed negative-binomial distribution of acrosomal proteomes performed on spermatozoa with short (upper panels) and long apical hooks (lower panels). Note the similar median values (red line).

### 2.2. Interspecific and intraspecific diversification of acrosomal proteomes

First, we aimed to find out whether acrosomes vary between rodent species at the protein level. As shown in Fig. 3A, the Sparse Partial Least-Squares Discriminant Analysis (sPLS-DA) clearly discriminated acrosomal content between species with MM having the most significantly differentiating proteome of the acrosome (for both Comp1 and Comp2: AUC = 1, p = 0.0007) regardless the sperm morphology. To identify proteins that contribute to interspecific differences in the acrosome composition, we extracted and plotted loading values of the sPLS-DA from Fig. 3A. Among the top 20 loadings of components 1 (Comp1) and 2 (Comp2) were identified proteins representing various biological processes and functions, Fig. 3D,G. The most represented functional groups included proteins essential for the process of fertilisation (e.g., SPACA3, TEX101, ZP3R, and ACRBP) as well as structural proteins of the cytoskeleton (e.g., ACTC1, VIM, GSN, and CAPG). In addition, the loading plots revealed proteins involved in biochemical processes such as lipid metabolism (e.g., GSTM1, ACOT7), tricarboxylic acid cycle (e.g., MDH1, PDHA1), glycolysis (e.g., GAPDHS) and fatty acid metabolism (e.g., ACOT7). Other notable proteins included members of the lipocalin protein family mediating chemical communication and sperm maturation (Janotová and Stopka, 2009, Stopková et al., 2021), namely LCN5, PTGDS and FABP9. These protein categories were mostly characterizing acrosomal content in species of MM or AF. In contrast, antioxidant enzymes (i.e., PRDX1, PRDX6) were common to MM and MA, while proteasomal subunits, chaperones and other regulators assisting protein folding and response to unfolded proteins (i.e., PSMA7, PDIA4, MANF) were exclusively prominent in MA acrosome.

**Fig. 3.**
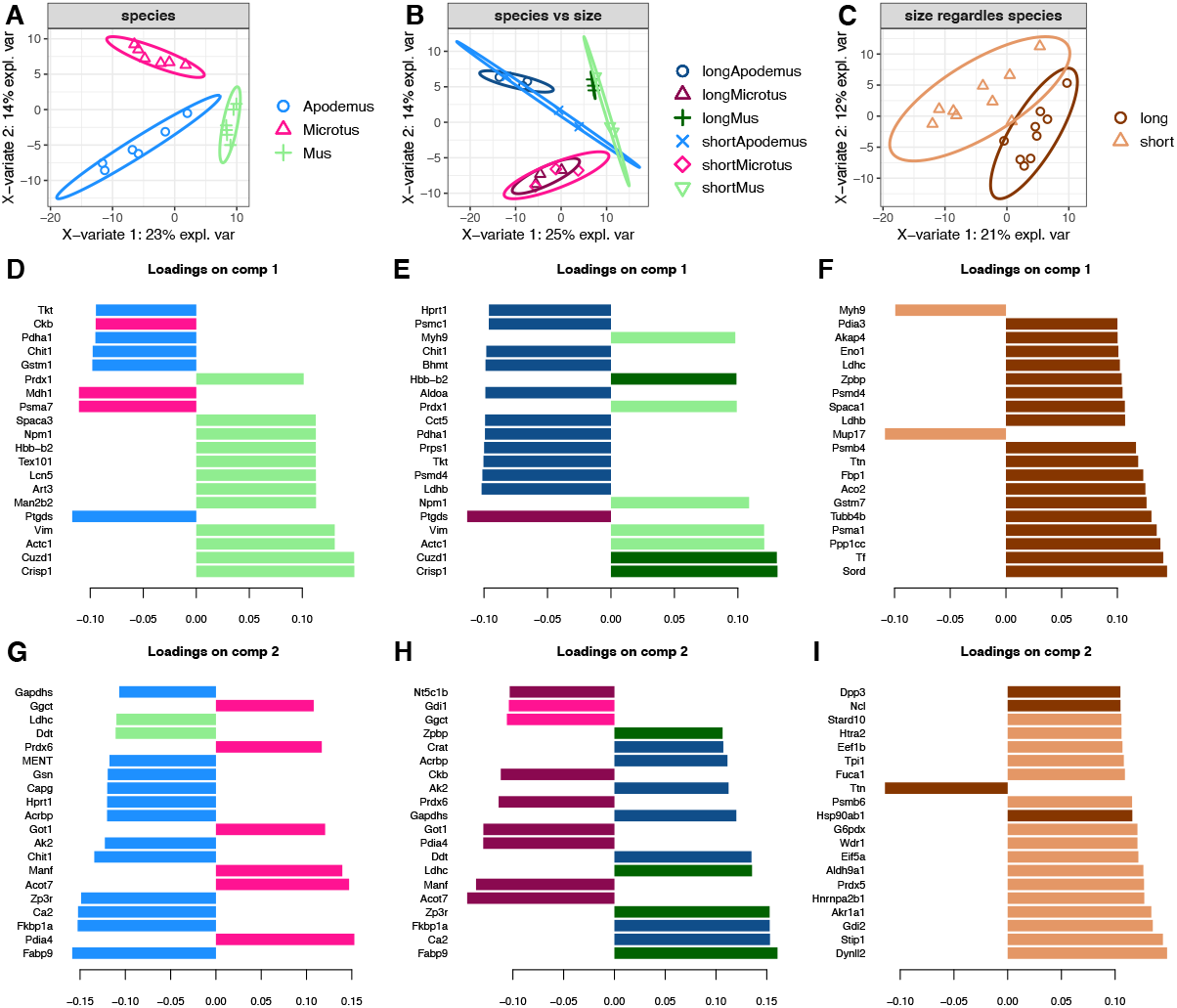
Diversification of acrosomal proteomes. (A-C) Sparse Partial Least Squares Discriminant Analysis (sPLS-DA) demonstrated three distinct species-biased clusters of acrosomal proteins (A), within-species proteomic variation regarding the length of apical hooks (B) and differentiation between short and long apical hook proteomes regardless the species (C). Note the highest and lowest intraspecific variation in species of *M. musculus* (green) and *M. arvalis* (pink), respectively. (D-I) Loading plots of the top 20 proteins contributing the most to differences between tested groups. Colors indicate the species or sample in which the protein is highly abundant. Proteins are ordered bottom-to-top as the most-to-least important in species separation.

Considering the length of the apical hooks, Fig. 3B, the discriminant analysis revealed the highest proteomic variation between individuals in MM. On the other side, the lowest intraspecific variation was observed in MA. Noteworthy, loading plots identified a group of the same proteins that contribute the most on Comp1 and Comp2 as in Fig. 3D,G, indicating a similar trend at the intraspecific level, see the bottom proteins in Fig. 3E,H. In addition to above mentioned ACTC1, VIM, PTGDS, FABP9, ZP3R and ACOT7, these included proteins involved in sperm maturation (i.e., CRISP1), cell-cell interaction (i.e., CUZD1) and protein folding (i.e., PDIA4). When selecting apical hook length regardless of species as a discriminative feature, sPLS-DA detected two different clusters representing short and long apical hook proteomes (for Comp2 AUC = 1, p = 0.0003), Fig. 3C. We observed a partial overlap of proteins responsible for morphological differences with those in Fig. 3E,H, namely PSMD4, ZPBP, MYH9, LDHB and LDHC that are involved in protein degradation, sperm-oocyte binding, regulation of actin dynamics and pyruvate metabolism, respectively, Fig. 3F,I.

### 2.3. Markers of short and long acrosomes

Based on the observed proteomic variation in acrosomal content with regard to length of apical hooks in Fig. 3, we employed Power Law Global Error Model (PLGEM) to identify significant and differentially abundant proteins (i.e., p < 0.05, fold difference abs(FD) > 2) differentiating short and long acrosomes within the species, Fig. 4A-C. Because such analysis does not estimate variable importance, we further applied a machine learning approach termed *Random forest*, Fig. 4D-F. This method uses for classification besides bootstrap also ‘out-of-bag’ samples and assigns each protein an importance value (here cut-off 0.03) in contribution to sample differentiation. Notably, the same functional protein categories as those separating species were again identified as responsible for individual variation. Although both these approaches revealed a similar trend of differentiation, *Random forest* analysis detected also those proteins that were with PLGEM identified as non-significant. As shown in Fig. 4, spermatozoa that possess short apical hooks were the best characterized by cytoskeletal proteins together with proteins involved in cell adhesion (MM-biased). On the contrary, fertilisation proteins (MM-biased), members of the ubiquitin-proteasome system (MM- and AF-biased) and proteins of the carbohydrate metabolic process (of importance in all three species) were detected in sperm with long apical hooks. Notably, several proteins were identified with all three applied analyses, i.e., sPLS-DA, PLGEM and *Random forest*, as separating acrosomal proteomes of short from long apical hooks. These included myosin-9 (MYH9), gamma-glutamylcyclotransferase (GGCT), protein similar to major urinary protein 17 (sMUP17), vimentin (VIM), actin alpha cardiac muscle 1 (ACTC1) significantly enriched in samples of short acrosomes, while serotransferrin (TF), sperm acrosome membrane-associated protein 1 (SPACA1), zona pellucida sperm-binding protein 3 receptor (ZP3R), and lactate dehydrogenase C chain (LDHC) were of significance in long acrosomes. Although the role of many of these proteins in the acrosomal content is still unknown, they can be considered as potential markers of apical hook length and thus fertility in muroid rodents.

**Fig. 4.**
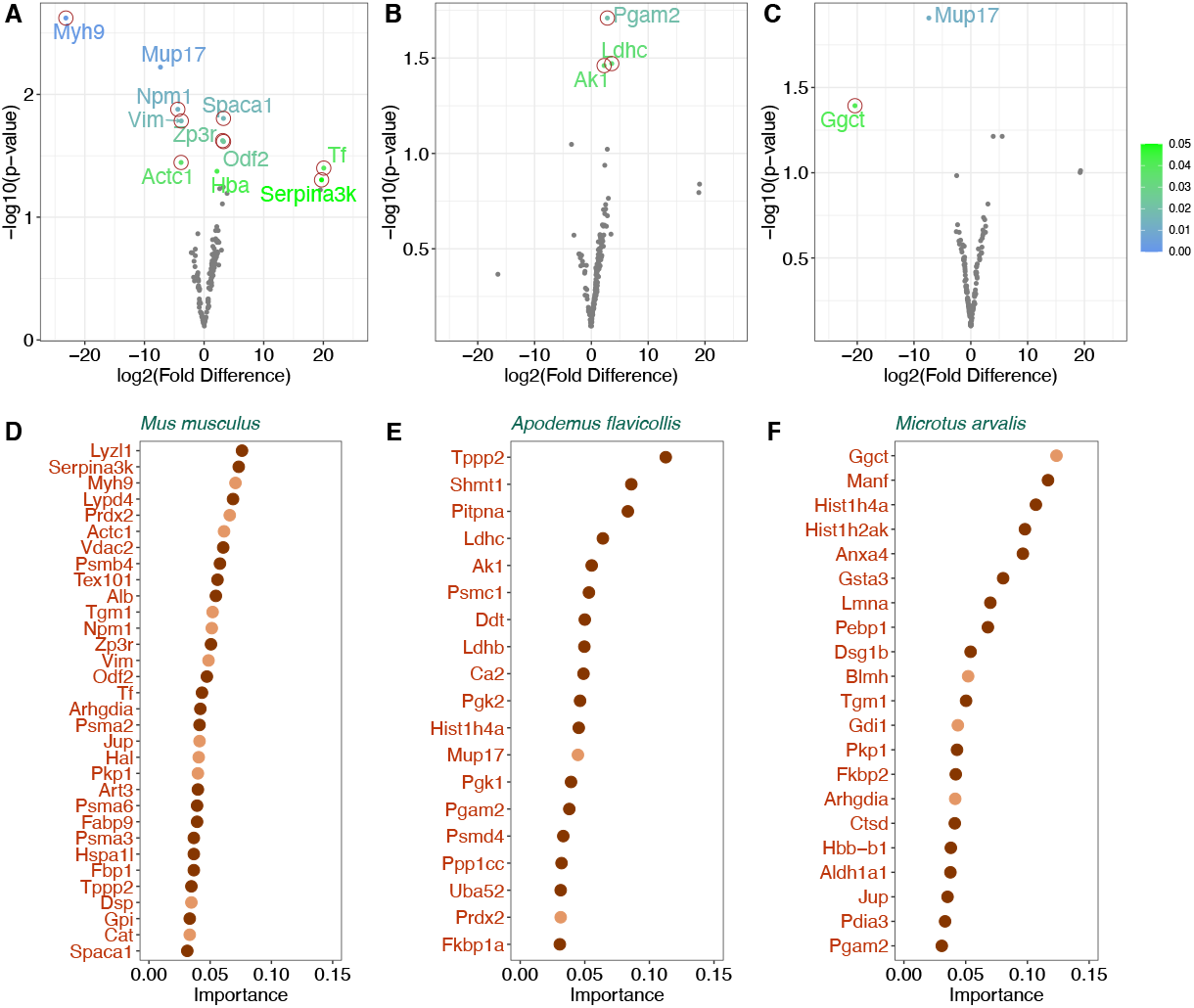
Markers of short and long acrosomes. (A-C) Volcano plots showing proteins differentially abundant (abs (FD > 2)) in sperm that possess short and long apical hooks (left and right from FD = 0, respectively). Significant proteins (p < 0.05) are labeled with gene names and color-coded based on their significance values (see the vertical bar). Red circles indicate proteins detected with *Random forest* in Fig. 4D-F. (D-F) Acrosomal proteins identified with *Random forest* algorithm as the best-differentiating spermatozoa with short (light brown) and long apical hooks (dark brown). Individual proteins are ordered top-to-bottom based on the highest-to-lowest importance in sperm separation by acrosome length. *Mus musculus* (A, D), *Apodemus flavicollis* (B, E), *Microtus arvalis* (C, F).

## 3. Discussion

Levels of sperm competition are associated with variation in the design, performance and molecular composition of spermatozoa across rodent species (Gómez Montoto et al., 2011, Tourmente et al., 2015b, delBarco-Trillo et al., 2015, Vicens et al., 2017). Here, we aimed to determine whether these differences are detectable between males of one species and how selective forces act on competing males within species. To correlate genotype with phenotype, we selected individuals for nLC-MS/MS analysis of acrosomal content based on sperm morphometry. As expected from previous studies (Immler et al., 2007, Šandera et al., 2013), the length of the apical hook was the most variable sperm trait with the highest values observed in highly promiscuous species of AF. Also, the fact that we identified the most proteins in AF acrosomal proteome can be likely explained by the risk of sperm competition. Comparable results were also obtained by Vicens et al. (Vicens et al., 2017) in their proteomic study performed on closely related *Mus* species.

According to sPLS-DA analysis, proteins involved in sperm-oocyte interaction during fertilisation, cytoskeleton dynamics and metabolic processes were found as important in contributing to interspecific proteomic variation. Furthermore, many of these proteins representing different cellular functions separated acrosomal content across species regarding the apical hook length. This suggests that there are protein categories responsible besides between-species differences also for between-male variation in sperm morphology. Since most of these proteins were identified in MM and AF that belong to family *Muridae*, our data further indicate that phylogeny, rather than sperm competition, might have driven the diversification of sperm size and shape in these phylogenetically related murine rodents. Nevertheless, further research is needed to address our hypothesis on a higher number of species.

Further evidence of a possible relationship between intraspecific variation in apical hook size and proteomic content of sperm acrosomes was revealed with the PLGEM model and *Random forest*. Although *Random forest* appears as more sensitive in the detection of a trend of proteins separating short and long acrosomes, many of them were not identified as significant with the PLGEM model. Thus, from a methodological point, we demonstrate the accuracy of using a combination of deep machine learning approach and conventional statistical methods to determine proteins that cause variation within a small sample size. Using three types of analyses, i.e., discriminant analysis, differential abundance and machine learning, several proteins were repeatedly detected and thus likely represent markers of apical hook length in examined species of murine rodents. As significant, highly abundant and best-differentiating samples of short apical hooks were identified proteins that mediate cytoskeleton organization and cell motility (MYH9, VIM, ACTC1, all detected in MM), apoptosis (GGCT, in MA) and chemical communication (sMUP17, in MM and MA, in AF nonsignificant). On the other side, proteins that play a role in binding and transport of iron (TF, detected in MM), acrosome dispersion and sperm-zona pellucida interactions (SPACA1, ZP3R, all in MM) and pyruvate fermentation (LDHC, in AF) characterized acrosomal content isolated from sperm with long hooks.

In line with the highest intraspecific proteomic variation observed in MM, the most functional protein categories separating spermatozoa with short and long apical hooks were found in MM. Cytoskeletal proteins together with lipocalin sMUP17 were highly abundant in short apical hooks. Although significance of sMUP7 and other detected lipocalins for male reproduction needs to be addressed, these findings are supportive of our recent study (Otčenášková et al., 2023). Enrichment of proteins involved in sperm-oocyte interactions including zona pellucida-binding, protein degradation and gluconeogenesis indicates that sperm with longer hooks are of better fertilisation competency and produce more energy as already postulated by (Varea-Sánchez et al., 2016). In line with this, serotransferrin TF was found positively correlated with sperm quality (Xin et al., 2019). Our data further support the evidence that glycolysis is rather utilized for energy production than oxidative phosphorylation in less promiscuous species such as MM (Tourmente et al., 2015a). Moreover, similar trend (i.e., high abundance of proteins of gluconeogenesis, proteasomal subunits and motility proteins in long acrosomes) was observed in highly promiscuous AF reflecting phylogenetic relationship of these species and partially also promiscuous mating strategy in AF. In samples isolated from MA were due to low between-male variation detected only two differentially abundant proteins being GGCT and sMUP17 significantly enriched in spermatozoa with short apical hooks.

From the evolutionary point of view, our data indicate that interspecific differences in sperm morphology and corresponding proteomic variation can also be detected on the within-species level. This suggests that there is an evolvability of the mechanisms of sperm competition that can be driven by selection at the regulatory level in species with different levels of promiscuity. In other way, this paper shows that there is innate variability in muroid rodents that can be reconfigured by selection when multi-male mating is more common. We are aware of limitations of this study, such as low number of species. However, a combination of several bioinformatics techniques clearly show that morphology variation can be detected on the level of proteomes thus revealing potential markers of selection at least in our selected rodent species. This study also has the potential to become an inspiration for other studies that would extend our findings in other species of mammals including humans.

## 4. Materials and Methods

### 4.1. Animals

Three species of muroid rodents, i.e., the house mouse (*Mus musculus musculus*), the yellow-necked mouse (*Apodemus flavicollis*) and the common vole (*Microtus arvalis*) were trapped in the field during the breeding season (May – July 2020) in locations near Vestec (49°58’50.2”N, 14°29’08.3”E; 49°59’32.3”N, 14°29’24.3”E) and Brandys nad Labem (50°11’12.874”N, 14°40’5.350”E), Czech Republic. From captured animals, only adult males (N = 31) were examined (eight to 14 individuals for each species), while juveniles and females were released back to the wild. Males were sacrificed by cervical dislocation on the day of capture and immediately weighed, followed by removal and weighing of the testes. All animal procedures strictly followed the laws of the Czech Republic, paragraph 17 no. 246/1992. This study was conducted with the approval of the local ethics committee of the Faculty of Science, Charles University.

### 4.2. Sample collection and sperm morphometry

Fresh sperm samples were recovered from distal portions of *cauda epididymis*. These were placed into two droplets of pre-tempered M2 fertilising medium (200 μL) (Sigma-Aldrich, M7167) covered with highly viscous paraffin oil (Carl Roth, 8904.1) and incubated at 37°C and 5% CO_2_, 10 min to allow sperm to swim out. The presence and viability of sperm were checked. An aliquot (5 μL) of isolated sperm was smeared onto a glass. Air-dried smears were fixed in 3.7% formaldehyde (vol./vol.) in phosphate-buffered saline (PBS) pH 7.4, 10 min and twice rinsed in PBS, 5 min. Following the washing steps, fixed smears were stained in Giemsa-Romanowski (Penta, 14460-11000), 10 min, RT and twice washed in dH_2_O, 5 min. Spermatozoa were viewed with a light microscope Olympus BX43 (Japan) in phase contrast and images were taken. The following five sperm dimensions were manually measured in open-source Fiji v1.52p software (Schindelin et al., 2012): head length (HL), head width (HW), apical hook length, midpiece length and tail length (length of the principal piece and end piece together). The sperm head area was calculated using the formula (HL/2) × (HW/2) × π, which is equal to half the area of the ellipse. Morphometric analysis was performed only in spermatozoa with normal appearance (a total of 50 sperm per male). A complete table with sperm dimension measurements is available from the corresponding author upon reasonable request.

### 4.3. Immunofluorescence staining and confocal microscopy

Freshly released spermatozoa (5 μL) were twice washed in PBS, smeared and air-dried onto a glass. Following the fixation in 3.7% formaldehyde in PBS, 10 min, RT, slides were twice washed in PBS, 5 min. Samples were then blocked with 5% bovine serum albumin (BSA; Sigma-Aldrich, A7906) (vol./vol.) in PBS for 1 h, RT and incubated with Alexa Fluor 568-conjugated peanut agglutinin (PNA; Life Technologies, L32458) diluted 1:500 in 1% BSA, 30 min, RT, to highlight the acrosome. After rinsing in PBS twice and in dH_2_O once, nuclei were counterstained with VECTASHIELD^®^ Antifade medium with DAPI (Vector laboratories, H-1200-10) and slides were sealed with coverslips. Images were taken by a Carl Zeiss LSM 880 NLO confocal microscope using ZEN 2.1 software and processed in open-source Fiji software (Schindelin et al., 2012).

### 4.4. Sperm capacitation and acrosome reaction

Spermatozoa underwent capacitation and acrosome reaction following the procedure in (Otčenášková et al., 2023). In brief, isolated spermatozoa were collected and transferred to a total of eight Petri dishes with new pre-tempered M2 medium droplets (100 μL, three droplets per dish) at a concentration of 5 ×10^6^ sperm per one mL. After 90 min capacitation at 37°C and 5% CO_2_, sperm samples were pooled in two 1.5 mL Eppendorf tubes (four Petri dishes per tube) and subjected to centrifugation, 5 min, 0.4 RCF, RT. After removing supernatants, pre-tempered sterile PBS (600 μL) was added and the tubes were centrifuged again to wash out the medium. Supernatants were discarded and resuspended pellets were divided into eight 1.5 mL Eppendorf tubes filled with pre-tempered sterile PBS (200 μL). Acrosome reaction was induced by calcium ionophore (added at a final concentration of 5 μM; Sigma-Aldrich, C7522) and spermatozoa were further incubated for 90 min, 37°C, 5% CO_2_, for acrosomal exocytosis to occur.

### 4.5. Protein isolation and proteomic analysis

Samples of spermatozoa that completed acrosome reaction were merged into one 1.5 mL Eppendorf tube per male and centrifuged, 14000 RCF, 10 min, 4°C, to separate acrosomal content and acrosome-reacted sperm. While supernatants were removed and kept on ice, sperm pellets were for 1 h, RT, suspended in rehydration buffer (100 μL) mixed from 8 M urea (Sigma-Aldrich, U6504), thiourea (Sigma-Aldrich, T8656), 1% Chaps (Sigma-Aldrich, C9426), 20 mM DTT (Sigma-Aldrich, D9163) and dH_2_O. Proteins in both supernatants and sperm pellets were precipitated with ice-cold acetone (added to samples in a 4:1 vol./vol. ratio), and samples were incubated overnight at -20°C. Following centrifugation on the next day, 14000 RCF, 10 min, 4°C, acetone was carefully discarded and prepared samples (N = 62, two samples per individual) were stored at -80°C until needed.

According to our morphometric analysis, we selected three individuals per species with the lowest and three individuals with the highest length of apical hook for nLC-MS/MS analysis. After samples of acrosomal proteins (N = 18) defrosted, they were suspended and digested in a buffer containing 1% SDC (Sigma-Aldrich, D6750) and 100 mM TEAB (Sigma-Aldrich, 86600), pH 8.5. The total protein content in lysates was quantified by the BCA assay kit (Thermo Fisher, 23227). Proteins (20 μg per sample) were reduced with 5 mM TCEP (a final concentration; Thermo Fischer, 20490), 1 h, 60°C, followed by the addition of blocking agent MMTS (10 mM; Thermo Fischer, 23011), 10 min, RT. Samples were subsequently digested with trypsin in 50:1 vol./vol. ratio, overnight at 37°C. Peptides were loaded onto the Michrom C18 column for desalting.

Desalted peptides were separated by nano-scale reversed phase liquid chromatography using EASY-Spray columns (50 cm × 75 μm ID, PepMap C18, 2 μm particles, 100 Å pore size). Mobile phases A and B were composed of 2% acetonitrile and 0.1% formic acid in water and 0.1% formic acid in 80% acetonitrile, respectively. Peptides were suspended in a loading buffer containing water, 2% acetonitrile and 0.1% trifluoroacetic acid, and loaded onto an Acclaim™ PepMap300 C18 trap column (300 μm × 5 mm, 5 μm, 300 Å pore size) at a flow rate of 15 μl min^-1^ for 4 min. The gradient of mobile phase B was as follows: 4% B for 4 min, 4–35% B in 60 min, 35-75% B in 61 min, hold for 8 min, 75-4% B in 70 min, and 4% B for 15 min. Following the electrospray ionization of liquid eluates, originated gas phase ions were analysed by a Thermo Orbitrap Fusion™ (Q-OT-qIT; Thermo Fisher, USA) mass spectrometer under conditions described previously (Kuntová et al., 2018, Otčenášková et al., 2023, Stopková et al., 2023).

### 4.6. Statistical analysis and bioinformatics

Raw MS data were using the Andromeda search engine quantified by MaxQuant software version 1.6.34 against the modified Uniprot *Mus musculus* database (dated June 2015, with 44,900 entries) that we downloaded to edit lipocalin sequences from the ENSEMBL database. The false discovery rate was set to 1% for all the identifications and the minimum peptide length to seven amino acids. The between-run variations in raw proteomic dataset were removed by label-free LFQ normalisation (Cox et al., 2014). We performed all following data manipulations, filtering, statistical analyses in R software (Crawley, 2007). All protein identifications that were identified but not quantified (zero abundances), all proteins containing less than 2 unique peptides, and all contaminants detected in MaxQuant output files (porcine trypsin, bovine albumins, human keratins etc.) were removed. Following this data reduction, Quantile normalisation from ‘normalizerDE’ package was performed (Willforss et al., 2019). Next, the Sparse Partial Least Squares Discriminant Analysis (sPLS-DA) within the ‘mixOmics’ package was employed for data exploration (Rohart et al., 2017). The package ‘mixtools’ (Gentleman et al., 2004) was used to detect a possible change in data distribution after second normalisation. Differentially expressed/ abundant proteins were identified using the Power Law Global Error Model - PLGEM (Pavelka et al., 2004) with functions plgem.fit and plgem-stn (Gentleman et al., 2004). The algorithm Random Forest for Classification (Breiman, 2001) was employed to identify proteins that differentiate sperm with short and long apical hooks. Finally, R software with ggplot2 and related packages (Wickham, 2016) was used to generate all plots and figures. Raw and LC-MS/MS tables are provided in Supplementary dataset 1.

## Acknowledgements

We are grateful who helped with the lab work at the initial phase of the project. We acknowledge Karel Harant and Pavel Talacko from Proteomics Core Facility, BIOCEV, Faculty of Science, Charles University, Prague for the proteomic and mass spectrometric measurements. We thank Eliška Macíčková from Imaging Methods Core Facility at BIOCEV, institution supported by the MEYS CR (LM2023050 Czech-BioImaging), for acquisition of confocal data presented in this paper.

## Data Availability Statement

The mass spectrometry proteomics data have been deposited to the ProteomeXchange Consortium via the PRIDE (Perez-Riverol et al., 2022) partner repository with the dataset identifier PXD043385 and 10.6019/PXD043385. Resulting tables are available as xlsx file also in PRIDE.

## Author contributions

T.O.: Investigation, Visualization, Writing – original draft, Writing – review & editing. T.D.: Investigation, Formal analysis. R.S.: Formal analysis, Writing – original draft, Writing – review & editing. P.S.: Conceptualization, Formal analysis, Resources, Visualization, Supervision, Funding acquisition, Writing – original draft, Writing – review & editing. All authors reviewed and approved the final version.

## References

Breed, W. G. 2004. The spermatozoon of Eurasian murine rodents: Its morphological diversity and evolution. Journal of Morphology, 261, 52–69.

Breiman, L. 2001. Random Forests. Machine Learning, 45, 5–32.

Briskie, J. V. & Montgomerie, R. 1992. Sperm size and sperm competition in birds. Proceedings. Biological sciences, 247, 89–95.

Clift, L. E., Andrlikova, P., Frolikova, M., Stopka, P., Bryja, J., Flanagan, B. F., Johnson, P. M. & Dvorakova-Hortova, K. 2009. Absence of spermatozoal CD46 protein expression and associated rapid acrosome reaction rate in striped field mice (Apodemus agrarius). Reproductive biology and endocrinology, 7, 29.

Cox, J., Hein, M. Y., Luber, C. A., Paron, I., Nagaraj, N. & Mann, M. 2014. Accurate proteome-wide label-free quantification by delayed normalization and maximal peptide ratio extraction, termed MaxLFQ. Molecular & cellular proteomics, 13, 2513–2526.

Crawley, M. J. 2007. The R Book, Chechester, Wiley Publishing.

Delbarco-Trillo, J., Mateo, R. & Roldan, E. R. 2015. Differences in the fatty-acid composition of rodent spermatozoa are associated to levels of sperm competition. Biology open, 4, 466–473.

Firman, R. C. & Simmons, L. W. 2008. The frequency of multiple paternity predicts variation in testes size among island populations of house mice. Journal of evolutionary biology, 21, 1524–1533.

Fisher, H. S. & Hoekstra, H. E. 2010. Competition drives cooperation among closely related sperm of deer mice. Nature, 463, 801–803.

Gentleman, R. C., Carey, V. J., Bates, D. M., Bolstad, B., Dettling, M., Dudoit, S., Ellis, B., Gautier, L., Ge, Y., Gentry, J., Hornik, K., Hothorn, T., Huber, W., Iacus, S., Irizarry, R., Leisch, F., Li, C., Maechler, M., Rossini, A. J., Sawitzki, G., Smith, C., Smyth, G., Tierney, L., Yang, J. Y. & Zhang, J. 2004. Bioconductor: open software development for computational biology and bioinformatics. Genome Biol, 5, R80.

GÓmez Montoto, L., Varea SÁnchez, M., Tourmente, M., MartÍN-Coello, J., Luque-Larena, J. J., Gomendio, M. & Roldan, E. R. 2011. Sperm competition differentially affects swimming velocity and size of spermatozoa from closely related muroid rodents: head first. Reproduction, 142, 819–830.

Hook, K. A., Wilke, L. M. & Fisher, H. S. 2021. Apical sperm hook morphology is linked to sperm swimming performance and sperm aggregation in Peromyscus mice. Cells, 10.

Immler, S., Moore, H. D., Breed, W. G. & Birkhead, T. R. 2007. By hook or by crook? Morphometry, competition and cooperation in rodent sperm. PloS one, 2, e170.

JanotovÁ, K. & Stopka, P. 2009. Mechanisms of chemical communication: the role of Major Urinary Proteins. Folia Zoologica, 58, 41–55.

Johnson, P. M., Clift, L. E., Andrlikova, P., Jursova, M., Flanagan, B. F., Cummerson, J. A., Stopka, P. & Dvorakova-Hortova, K. 2007. Rapid sperm acrosome reaction in the absence of acrosomal CD46 expression in promiscuous field mice (Apodemus). Reproduction, 134, 739–747.

KuntovÁ, B., StopkovÁ, R. & Stopka, P. 2018. Transcriptomic and proteomic profiling revealed high proportions of odorant binding and antimicrobial defense proteins in olfactory tissues of the house mouse. Frontiers in genetics, 9, 26.

Moore, H., DvorÁKovÁ, K., Jenkins, N. & Breed, W. 2002. Exceptional sperm cooperation in the wood mouse. Nature, 418, 174–177.

OtčenÁŠkovÁ, T., MacÍčkovÁ, E., VondrÁkovÁ, J., FrolÍkovÁ, M., Komrskova, K., StopkovÁ, R. & Stopka, P. 2023. Proteomic analysis of the mouse sperm acrosome - towards an understanding of an organelle with diverse functionality. European journal of cell biology, 102, 151296.

Parker, G. A. 1970. Sperm competition and its evolutionary consequences in the insects. Biological Reviews, 45, 525–567.

Pavelka, N., Pelizzola, M., Vizzardelli, C., Capozzoli, M., Splendiani, A., Granucci, F. & Ricciardi-Castagnoli, P. 2004. A power law global error model for the identification of differentially expressed genes in microarray data. BMC Bioinformatics, 5, 203.

Perez-Riverol, Y., Bai, J., Bandla, C., GarcÍA-Seisdedos, D., Hewapathirana, S., Kamatchinathan, S., Kundu, D. J., Prakash, A., Frericks-Zipper, A., Eisenacher, M., Walzer, M., Wang, S., Brazma, A. & VizcaÍno, J. A. 2022. The PRIDE database resources in 2022: a hub for mass spectrometry-based proteomics evidences. Nucleic acids research, 50, D543–D552.

Rohart, F., Gautier, B., Singh, A. & LÊ Cao, K.-A. 2017. mixOmics: An R package for ‘omics feature selection and multiple data integration. PLOS Computational Biology, 13, e1005752.

Roldan, E. R., Gomendio, M. & Vitullo, A. D. 1992. The evolution of eutherian spermatozoa and underlying selective forces: female selection and sperm competition. Biological reviews of the Cambridge Philosophical Society, 67, 551–593.

Rowe, M., Laskemoen, T., Johnsen, A. & Lifjeld, J. T. 2013. Evolution of sperm structure and energetics in passerine birds. Proceedings. Biological sciences, 280, 20122616.

Šandera, M., Albrecht, T. & Stopka, P. 2013. Variation in apical hook length reflects the intensity of sperm competition in murine rodents. PloS one, 8, e68427.

Schindelin, J., Arganda-Carreras, I., Frise, E., Kaynig, V., Longair, M., Pietzsch, T., Preibisch, S., Rueden, C., Saalfeld, S., Schmid, B., Tinevez, J. Y., White, D. J., Hartenstein, V., Eliceiri, K., Tomancak, P. & Cardona, A. 2012. Fiji: an open-source platform for biological-image analysis. Nature methods, 9, 676–682.

Stockley, P., Gage, M. J., Parker, G. A. & MØller, A. P. 1997. Sperm competition in fishes: the evolution of testis size and ejaculate characteristics. The American naturalist, 149, 933–954.

StopkovÁ, R., MatĚjkovÁ, T., DodokovÁ, A., Talacko, P., Zacek, P., Sedlacek, R., PiÁkek, J. & Stopka, P. 2023. Variation in mouse chemical signals is genetically controlled and environmentally modulated. Scientific reports, 13, 8573.

StopkovÁ, R., OtčenÁŠjovÁ, T., MatĚjkovÁ, T., KuntovÁ, B. & Stopka, P. 2021. Biological roles of lipocalins in chemical communication, reproduction, and regulation of microbiota. Frontiers in physiology, 12, 740006.

StØstad, H. N., Johnsen, A., Lifjeld, J. T. & Rowe, M. 2018. Sperm head morphology is associated with sperm swimming speed: A comparative study of songbirds using electron microscopy. Evolution; international journal of organic evolution, 72, 1918–1932.

Tourmente, M., Gomendio, M. & Roldan, E. R. 2011. Sperm competition and the evolution of sperm design in mammals. BMC evolutionary biology, 11, 12.

Tourmente, M., Villar-Moya, P., Rial, E. & Roldan, E. R. 2015a. Differences in ATP generation via glycolysis and oxidative phosphorylation and relationships with sperm motility in mouse species. The Journal of biological chemistry, 290, 20613–20626.

Tourmente, M., Villar-Moya, P., Varea-SÁnchez, M., Luque-Larena, J. J., Rial, E. & Roldan, E. R. 2015b. Performance of rodent spermatozoa over time is enhanced by increased ATP concentrations: the role of sperm competition. Biology of reproduction, 93.

Tourmente, M., Zarka-Trigo, D. & Roldan, E. R. 2016. Is the hook of muroid rodent’s sperm related to sperm train formation? Journal of evolutionary biology, 29, 1168–1177.

Varea-SÁnchez, M., Tourmente, M., Bastir, M. & Roldan, E. R. 2016. Unraveling the Sperm Bauplan: Relationships Between Sperm Head Morphology and Sperm Function in Rodents. Biology of reproduction, 95, 25.

Vicens, A., Borziak, K., Karr, T. L., Roldan, E. R. S. & Dorus, S. 2017. Comparative sperm proteomics in mouse species with divergent mating systems. Molecular biology and evolution, 34, 1403–1416.

Wickham, H. 2016. ggplot2: Elegant Graphics for Data Analysis., New York, Springer-Verlag.

Willforss, J., Chawade, A. & Levander, F. 2019. NormalyzerDE: Online Tool for Improved Normalization of Omics Expression Data and High-Sensitivity Differential Expression Analysis. Journal of proteome research, 18, 732–740

Xin, M., Vechtova, P., Shaliutina-Kolesova, A., Fussy, Z., Loginov, D., Dzyuba, B., Linhart, O., Boryshpolets, S., Rodina, M., Li, P., Loginova, Y. & Sterba, J. 2019. Transferrin identification in sterlet (Acipenser ruthenus) reproductive system. Animals (Basel), 9, 753.

